# Leptin Signalling in the Ovary of Diet-Induced Obese Mice Regulates Activation of Nod-Like Receptor Protein 3 Inflammasome

**DOI:** 10.1101/2021.08.16.456479

**Authors:** Marek Adamowski, Karolina Wołodko, Joana Oliveira, Juan Castillo-Fernandez, Daniel Murta, Gavin Kelsey, António M. Galvão

**Affiliations:** Department of Reproductive Immunology and Pathology, Institute of Animal Reproduction and Food Research of Polish Academy of Sciences, Olsztyn, Poland; Centro de Investigação em Ciências Veterinárias, Lusófona University, Lisbon, Portugal; Epigenetics Programme, The Babraham Institute, Cambridge, CB22 3AT, UK; Centro de Investigação Interdisciplinar Egas Moniz (CiiEM), Escola Superior de Saúde Egas Moniz, Campus Universitário, Quinta da Granja, Monte de Caparica, Portugal; C.I.I.S.A., Faculty of Veterinary Medicine, University of Lisbon, Lisbon, Portugal; Centre for Trophoblast Research, University of Cambridge, Cambridge, CB2 3EG, UK

**Keywords:** ovary, inflammation, leptin, obesity, NLRP3inflammasome

## Abstract

Obesity leads to ovarian dysfunction and the establishment of local leptin resistance. The aim of our study was to characterise levels of Nod-Like Receptor Protein 3 (NLRP3) inflammasome activation during obesity progression in the mouse ovaries and liver and test the putative role of leptin on its regulation. C57BL/6J mice were treated with equine chorionic gonadotropin (eCG) or human chorionic gonadotropin (hCG) for oestrous cycle synchronisation and ovaries collection. In diet-induced obesity (DIO) model, mice were fed chow diet (CD) or high fat diet (HFD) for 4 or 16 weeks (wk), whereas in hyperleptinemic model (LEPT), mice were injected with leptin for 16 days (16L) or saline (16C) and in the genetic obese leptin-deficient *ob/ob* (+/? and -/-) animals were fed CD for 4wk. Either ovaries and liver were collected, as well as cumulus cells (CCs) after superovulation from DIO and LEPT. In DIO protocol, protein expression of NLRP3 inflammasome components was increased in 4wk HFD, but decreased in 16wk HFD. Moreover LEPT and *ob/ob* models revealed NLRP3 and IL-1β upregulation in 16L and downregulation in *ob/ob.* Transcriptome analysis of CC showed common genes between LEPT and 4wk HFD modulating NLRP3 inflammasome. Moreover analysis in the liver showed upregulation of NLRP3 protein only after 16wk HFD, but also the downregulation of NLRP3 protein in *ob/ob-/-*. We showed the link between leptin signalling and NLRP3 inflammasome activation in the ovary throughout obesity progression in mice, elucidating the molecular mechanisms underpinning ovarian failure in maternal obesity.

## 2 Introduction

Obesity leads to self-directed tissue inflammation, a process mostly promoted by the continuous expansion of adipose tissue (Hotamisligil and Erbay 2008; Odegaard and Chawla 2008). Furthermore, literature presents a solid link between obesity and reproductive failure in women (Chu et al. 2007). Indeed, obesity in women has been associated to ovarian dysfunction, embryo implantation failure, abortion, foetal congenital abnormalities, and adult offspring adiposity and metabolic dysfunction (Chu et al. 2007; Penzias 2012; Robker 2008; Samuelsson et al. 2008). The ovaries from mice fed high fat diet (HFD) showed increased apoptosis and fewer mature oocytes (Jungheim et al. 2010). Furthermore, due to lipid accumulation, endoplasmic reticulum (ER) stress, mitochondrial dysfunction and increased ovarian cell apoptosis, these mice displayed anovulation and reduced *in vivo* fertilization rates (Wu et al. 2010), as well as abnormal embryo development (Minge et al. 2008). We have recently demonstrated the establishment of leptin resistance in the ovaries of mice treated with HFD (Wołodko et al. 2020) was mostly mediated by suppressor of cytokine signalling 3 (SOCS3). Hence, changes in local leptin signalling were shown to contribute to the pathophysiology of ovarian failure in obese females (Wołodko et al. 2021).

The inflammasome is a large intracellular protein complex that contains a cytosolic pattern recognition receptor. Among NOD-like receptors (NLR), the NLR protein 3 (NLRP3) inflammasome has been best characterised as a complex of proteins responsible for controlling the activity of two proinflammatory cytokines interleukin (IL)-1β and IL-18 (Davis, Wen, and Ting 2011; De Nardo and Latz 2011; Martinon, Burns, and Tschopp 2002). Activation of the pattern recognition receptor NLRP3 can be accomplished through two major signals: (i) priming signal, induced by the toll-like receptor (TLR)/nuclear factor (NF)-κB pathway; and (ii) pathogen-associated molecular patterns (PAMPs) and damage-associated molecular patterns (DAMPs) leading to assembly of inflammasome (Lamkanfi and Dixit 2014; Martinon, Burns, and Tschopp 2002). Both mechanisms lead to the recruitment of the adapter apoptosis-associated speck-like protein containing a C-terminal caspase recruitment domain (ASC), resulting in the activation of pro-caspase-1 (CASP1) and cleavage into the active form (Davis, Wen, and Ting 2011). The formation and activation of the inflammasome is possible through ASC, which links NLRP3 to CASP1 by means of its pyrin and caspase recruitment domain motifs (Martinon et al. 2006). Finally, activated CASP1 is known to process the maturation of IL-1β and IL-18 into active cytokines (Lamkanfi 2011). Importantly, obesity and insulin resistance (IR) have been associated with inflammation and subsequent activation of NLRP3 inflammasome (Traba and Sack 2017). The onset of inflammasome activation was also shown to be mediated by factors like glucose, ceramide, uric acid, or lipopolysaccharide (LPS) (Stienstra et al. 2012; Traba and Sack 2017). Furthermore, secondary signals could also come from extracellular ATP inducing K+ efflux; DAMPs/PAMPS leading to reactive oxygen species (ROS) production (Shao et al. 2015; Tözsér and Benkö 2016). Saturated free fatty acids (FFAs) were equally linked to inflammasome activation through both signals (Wen et al. 2011), as increased levels of FFAs are a general feature of obesity, IR or type-2 diabetes (Boden 2002; Krebs and Roden 2005). More recently, a link has been also established between NLRP3 inflammasome activation and levels of leptin signalling in various cellular contexts (Fu et al. 2017), corroborating the proinflammatory role of leptin (Cauble et al. 2018).

A recent report has shown the presence of NLRP3 inflammasome components at ovarian level during follicular development in mice, suggesting its involvement in ovulation (Z. Zhang, Wang, and Zhang 2019). Most importantly, NLRP3 was also suggested to be involved in the pathophysiology of polycystic ovary syndrome (PCOS) (Rostamtabar et al. 2020). Therefore, we presently hypothesise the regulation of NLRP3 in the ovary of obese mice is mediated by leptin signalling. We firstly confirmed NLRP3 inflammasome expression profile changed in the ovaries of cyclic mice. Subsequently, we also confirmed NLRP3 inflammasome components were differently expressed in the ovaries of 4 and 16 weeks (wk) diet induced obese (DIO) mice. Furthermore, using a mouse model of pharmacological hyperleptinemia (LEPT) and a genetic obese mouse which lacks leptin (*ob/ob*) we demonstrated the association between levels of leptin signalling and NLRP3 inflammasome activation in the ovary of obese mice. Moreover, we analysed the transcriptome of cumulus cells (CCs), the somatic companions of the oocyte, and concluded once more that leptin treatment-upregulated genes associated with NLRP3 inflammasome. Finally, we studied the NLRP3 inflammasome expression in the liver of DIO, LEPT and *ob/ob* mice and despite observing a different temporal signature in DIO, with regard to the ovary, we also found a consistent downregulation in NLRP3 inflammasome activity in *ob/ob*, which are obese and lack leptin.

## 3 Materials & Methods

### 3.1 Animals

Female B6 mice (8 wk old) and B6.Cg-Lepob/J (*ob/ob*) were housed in the Animal Facility of Institute of Animal Reproduction and Food Research, Polish Academy of Sciences in Olsztyn. Breeding pairs were purchased from Jackson Laboratories (Bar Harbor, ME). Mice were housed with free access to food and water for the duration of the study (humidity 50±10%; 23°C; 12L:12D cycle). All procedures were approved by the Local Animal Care and Use Committee for University of Warmia and Mazury, Olsztyn. Guidelines for animal experiments followed EU Directive 2010/63/EU. Throughout the whole experiments, mice were monitored for any sings of welfare or disease. At 21 days of age female progeny were weaned and housed in groups of 3–5 in plastic cages with fresh sawdust bedding. By 8 wk of age, one group of B6 mice was subjected to hormonal protocol, while the other group was segregated into two different dietary protocols matched for similar body weight. In DIO model (n=10/group) mice were placed on standard CD (Picolab Rodent diet 20, #5053) with 13% of calories coming from fat, or on HFD with 59 % of calories coming from fat (AIN-76A with 33% hydrogenated coconut oil; LabDiet) for 4 or 16 wk. Hyperleptinemia model (n=8/group) was utilized to mimic high level of leptin through its intraperitoneal injections twice a day at total dosage 100 µg/day (injected at 09:00 and 21:00), while the control group received saline injections(Recombinant Mouse Leptin, GFM26, Cell Guidance Systems). Regarding *ob/ob* model (n=6/group), mice were kept on CD until 12 wk of age.

### 3.2 Induction of oestrus and dioestrus stages

The oestrous cycle was monitored studying vaginal cytology. Cells collected via saline lavage were placed on glass slide and stained with Diff Quik® kit (Medion Diagnostics AG, Switzerland, DQ-ST). Oestrus was characterised by cornified epithelium cells; metoestrus by both cornified cells and leukocytes; dioestrus by predominant leukocytes; and pro-oestrus by nucleated cells, as previously described (Kyrönlahti et al. 2011).

Group of female B6 mice (8 wk old) was injected in oestrus stage with eCG (G4877, 5IU, Sigma Aldrich, Saint Louis, Missouri, USA) followed after 48 h by hCG (Chorulon, 5IU, MSD Animal Health, Boxmeer, Netherlands) and tissues collected 18-20 h later in E. The second group of mice were injected with hCG and tissues were collected 16-18 h later in D. To reduce variation between groups ovaries from females from the remaining experiments were collected in dioestrus stage.

### 3.3 Protein extraction and Western blotting analysis

Protein expression in mouse ovary and liver was assessed by Western blotting. Ovaries and livers were homogenized with RIPA buffer (R0278; Sigma) containing protease inhibitors (phenylmethylsulfonyl fluoride, PMSF and Protease Inhibitor Cocktail, P8340; Sigma-Aldrich, St. Louis, MO, US) and phosphatase inhibitors (Pierce Phosphatase Inhibitor Mini Tablets 88667; Thermo Fisher Scientific) and incubated on ice for 1 h while vortexing in the meantime. After centrifugation (20 000 g, 15 min, 4°C) the supernatants were collected and protein concentration was determined with the Smith (Smith 1985) copper/bicinchoninic assay (Copper (II) Sulfate, C2284; Sigma and Bicinchoninic Acid Solution, B9643, Sigma). Samples were run (40 µg of protein) on 10-18% polyacrylamide gels. Immunoblotting was performed using the following primary antibodies NLRP3 (AG-20B-0014-C100; Adipogen), CASP1 (ab108362; Abcam), IL-18 (ab71495; Abcam), β-actin (A2228; Sigma), glyceraldehyde 3-phosphate dehydrogenase (GAPDH, ab9485; Abcam) and then transferred to nitrocellulose (10600009; GE Healthcare Life Science) or polyvinylidene fluoride (PVDF) membrane (IPVH00010; Merck Millipore). The membranes were blocked in phosphatase buffered saline (PBS) solution containing 3% powdered milk for 1 h. Primary antibodies were used at 1:1 000 (NLRP3, CASP1) and 1:250 (IL-18) dilution and incubated overnight at 4°C. The following day, proteins were detected by incubating the membranes with a polyclonal anti-mouse horseradish peroxidase (HRP)-conjugated secondary (1:10 000, 31430; Thermo Fisher Scientific), polyclonal anti-rabbit HRP-conjugated secondary (1:20 000, 31460; Thermo Fisher Scientific), polyclonal anti-mouse alkaline phosphatase-conjugated secondary (1:10 000, 31321; Thermo Fisher Scientific) and polyclonal anti-rabbit alkaline phosphatase-conjugated secondary (1:10 000, A3687, Sigma) antibodies, for 1,5 h in chemiluminescence method or 2,5 h in colorimetric method at room temperature (RT). All antibodies specifications are summarised in **Table 1**. Immunocomplexes were visualized subsequently using chemiluminescence detection reagent (SuperSignal West Femto kit, 34095; Thermo Fisher Scientific) or chromogenic substrate NBT/BCIP diluted 1:50 (11681451001; Roche) in alkaline phosphate buffer. Band density for each of the target protein was normalised against β-actin for NLRP3 and IL-18, while GAPDH was used for CASP1 as a reference protein. Finally, bands were quantified using the ChemiDoc or VersaDoc MP 4000 imaging system (Bio-Rad). Quantitative measurements of blot intensity were performed using ImageLab software.

**Table 1.** Specification of antibodies used for Western blotting.

| Antibody name and specificity | Company, Cat no, RRID no | Antibody dilution |
| --- | --- | --- |
| Mouse monoclonal against NLR family pyrin domain containing 3 (NLRP3) | AdipoGen Cat# AG-20B-0014, RRID:AB_2490202 | 1:1000 |
| Rabbit monoclonal against Caspase 1 (CASP1) | Abcam Cat# ab108362, RRID:AB_10858984 | 1:1000 |
| Rabbit polyclonal against interleukin-18 (IL-18) | Abcam Cat# ab71495, RRID:AB_1209302 | 1:250 |
| Mouse monoclonal against β-actin | Sigma-Aldrich Cat# A2228, RRID:AB_476697 | 1:10 000 |
| Rabbit polyclonal against glyceraldehyde 3-phosphate dehydrogenase (GAPDH) | Abcam Cat# ab9485, RRID:AB_307275 | 1:2500 |
| Goat anti-Mouse IgG (H+L) Secondary Antibody, HRP | Thermo Fisher Scientific Cat# 31430, RRID:AB_228307 | 1:1000 |
| Goat anti-Rabbit IgG (H+L) Secondary Antibody, HRP | Thermo Fisher Scientific Cat# 31460, RRID:AB_228341 | 1:20 000 |
| Goat anti-Mouse IgG (H+L) Secondary Antibody, AP | Thermo Fisher Scientific Cat# 31321, RRID:AB_10959407 | 1:1000 |
| Goat Anti-Rabbit IgG (whole molecule) Secondary Antibody, AP | Sigma-Aldrich Cat# A3687, RRID:AB_258103 | 1:10 000 |

### 3.4 Total RNA Isolation and cDNA Synthesis

Total RNA was extracted from whole ovary and 10 mg of liver, using TRI reagent (T9424; Sigma Aldrich) following the manufacturer’s instructions. RNA samples were stored at -80°C. Concentration and quality of RNA was determined spectrophotometrically and the ratio of absorbance at 260 and 280 (A_260/280_) was analysed confirming good RNA quality. Subsequently, 2 μg of RNA was reverse transcribed into cDNA using Maxima First Strand cDNA Synthesis Kit for RT-qPCR (K1642; ThermoFisher Scientific) (Galvão et al. 2012).

### 3.5 Real-time PCR

Real-time PCR assays were performed in a 7900 Real-time System (Applied Biosystems), using a default thermocycler program for all genes: a 10 min preincubation at 95°C was followed by 45 cycles of 15 sec at 95°C and 1 min at 60°C. A further dissociation step (15 sec at 95°C, 15 sec at 60°C, and 15 sec at 95°C) ensured the presence of a single product. *Ribosomal protein L37 (Rpl37)* was chosen as a house keeping gene and quantified in each real-time assay together with target gene. Based on gene sequences in GenBank (National Center for Biotechnology Information), the primers for *Nlrp3, Casp1, Il-1β, Il-18, Asc, Il-10, Tnf*, which sequences are presented in **Table 2**, were designed using Primer Express 3.0 software (Applied Biosystems). All reactions were carried out in duplicates in 384-well plate (4309849; Applied Biosystems) in 12 µl of total solution volume (Galvão et al. 2014). The data were analysed using the real-time PCR Miner algorithm (S. Zhao and Fernald 2005).

**Table 2.**
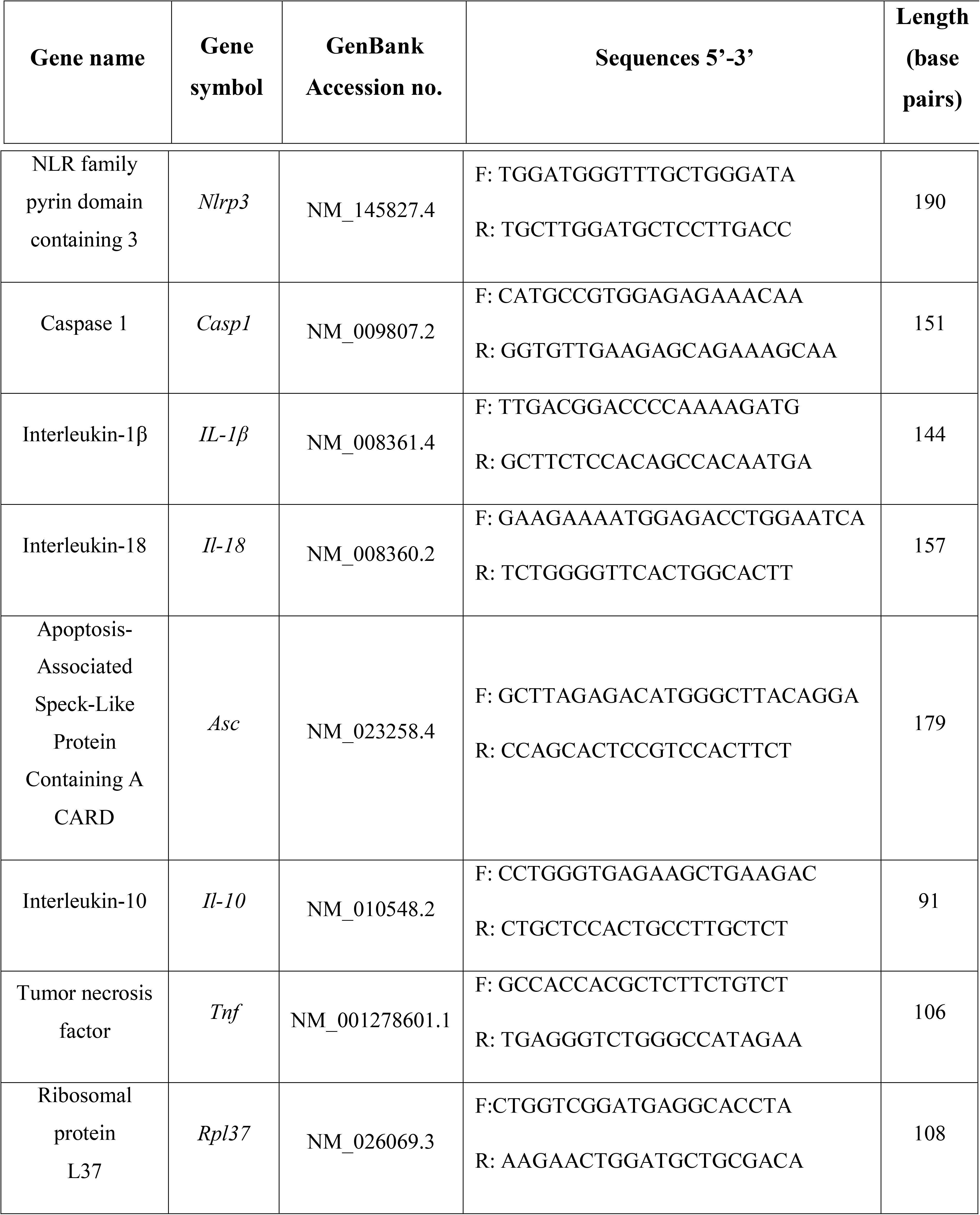
Specific primers used for quantitative real-time PCR.

### 3.6 ELISA immunoassay

The concentrations of IL-1β in tissue extracts of ovaries and livers were determined using an IL-1 beta Pro-form Mouse Uncoated ELISA kit (88-8014-22; Thermo Fisher Scientific) following manufacturer’s instructions. The standard curve concentrations ranged from 25 ng/ml to 3000 ng/ml and interassay coefficient variation (CV) was 7.27%.

### 3.7 Statistical analysis and data presentation

Statistical analyses were performed using the GraphPad Prism Software (Version 7.01, GraphPad Software, Inc.; La Jolla, CA, USA). Paired t test was employed to compare the changes of protein expression in mouse livers and ovaries. Comparisons of gene expression were performed using Wilcoxon matched pairs test. Results are presented as mean with standard deviation. Differences between the means for all tests were considered statistically significant if p< 0.05.

### 3.8 RNA-seq data from CCs

Methods followed the previously described (Wołodko et al. 2020). The GEO accession number for the dataset Sequencing is under submission.

## 4 Results

### 4.1 NLRP3 inflammasome components expression change in the ovary of cyclic mice

We first sought to characterise the expression of NLRP3-induced inflammasome components in the ovaries of mice, throughout the oestrous cycle. Fifteen female 8 wk old C57BL/6 (B6) mice were treated with hormones in order to synchronise oestrous cycle (**Figure 1A**). Mice received either equine chorionic gonadotropin (eCG) or human chorionic gonadotropin (hCG), as previously described (Hasegawa et al. 2016). Ovaries were collected in oestrus (E) and dioestrus (D) stage and further processed for mRNA or protein expression analysis, respectively. Real-time PCR analysis (n=7/group) revealed increased levels of *Casp1, Il-1*β and *Il-18* mRNA in D stage (**Figure 1B**, p<0.05). Moreover, Western blotting revealed increased NLRP3 protein expression in D (**Figure 1C**, p<0.05), as well as the pro-peptide (p24) and mature form (p18) of IL-18 (**Figure 1F, G**; p<0.01). Regarding CASP1, the long form (p45) was decreased in D (**Figure 1D**; p<0.05), but no significant changes were observed for the active CASP1 (p20) (**Figure 1E**). These results suggest the activation of NLRP3 inflammasome in D stage, through upregulation of NLRP3 and its downstream mediator IL-18. Next, we characterised the cellular distribution of NLRP3 protein in the ovaries collected from mice in D, using immunohistochemistry (IHC) and immunofluorescence (IF). We confirmed that NLRP3 protein was globally distributed in the ovary (**Figure 1I**). On the other hand, a closer observation with IHC revealed staining in granulosa cells (GC) and theca cells (TC), as well as oocytes, in all developmental stages of follicles in the ovary (**Figure 1K-M**). Negative controls stained with secondary antibodies did not reveal any brown staining (**Figure 1H, J**). The specificity of our IHC staining was corroborated by IF, in which a clear brown staining was observed in GC, TC and oocytes (**Figure 1O-U**). Negative control stained with rabbit immunoglobulin type G (IgG) confirmed no staining (**Figure 1N**). Our results once more corroborate the findings of Zhang and co-workers who not only observed the presence of NLRP3 protein in GC, TC and oocytes by IHC, but also confirmed the upregulation of NLRP3 in the ovaries of eCG treated mice (Z. Zhang, Wang, and Zhang 2019). Therefore, in subsequent experiments, collections were consistently performed in D.

**Figure 1.**
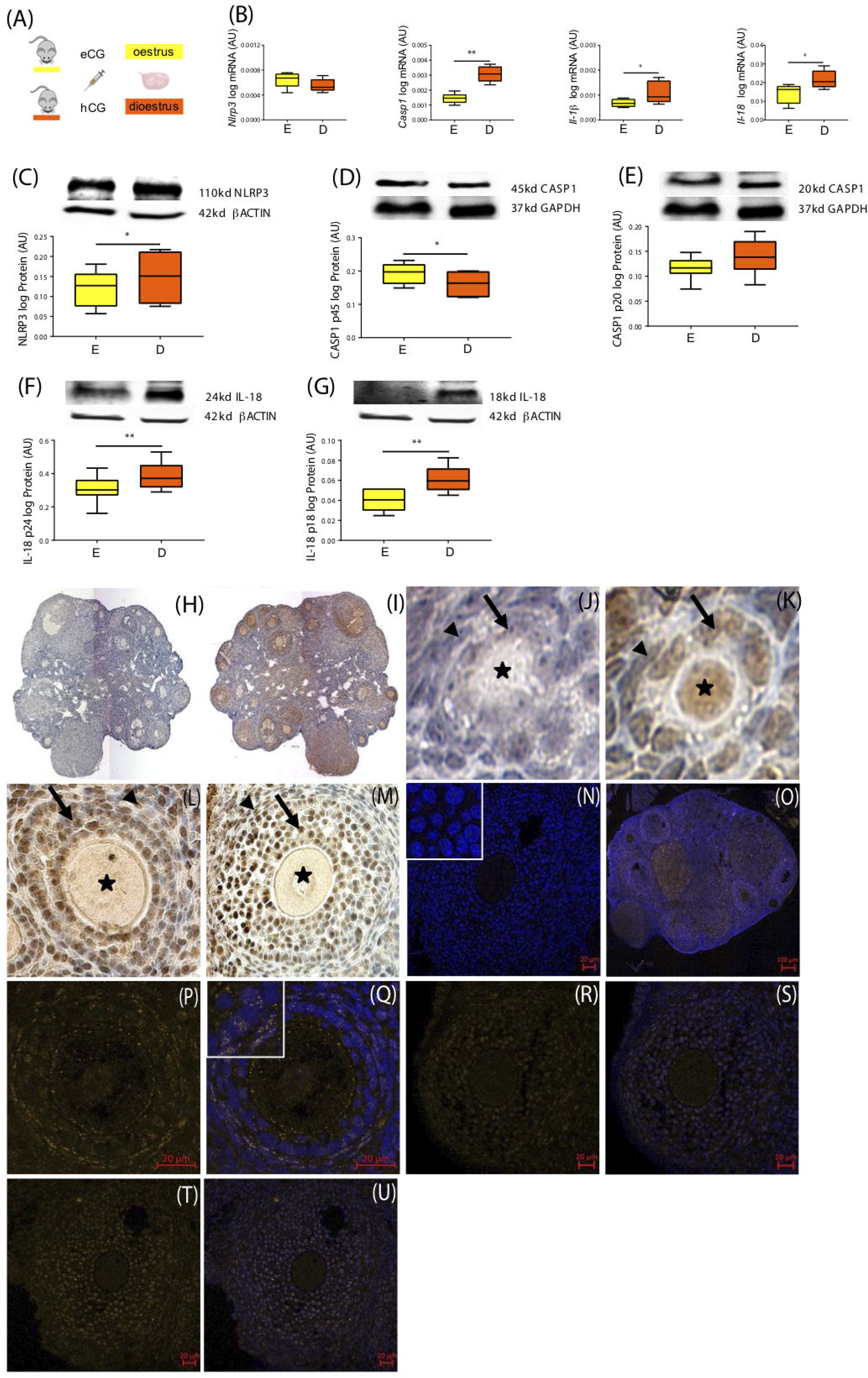
Morphofunctional characterisation of NLRP3 in the ovary of cyclic mice. (A) Experimental design: oestrous cycle synchronisation with eCG and hCG as previously described (Hasegawa et al. 2016). Ovaries were collected from animals in oestrus (E) or dioestrus (D) stage of the cycle. Quantification of mRNA levels of (B) Nod-Like Receptor Protein 3 (*Nlrp3*), caspase-1 (*Casp1*), interleukin-1β (*Il-1*β), interleukin-18 (*Il-18*) by real-time PCR. Abundance of (C) Nod-Like Receptor Protein 3 (NLRP3), (D) pro IL-18 p24, (E) IL-18 p18, (F) pro CASP1 p45 and (G) CASP1 p20 protein during E and D measured by western blotting analysis. Data was normalized to ribosomal protein L37 (*Rpl37*) mRNA expression and βactin of or glyceraldehyde 3-phosphate dehydrogenase (GAPDH) protein expression. Bars represent mean ± SEM. Statistical analysis between groups was carried out using Mann–Whitney. n=5-7 for real-time PCR analysis and n=6 for immunoblots. Asterisks indicate significant differences (*p< 0.05; **p<0.01). Representative immunohistochemical staining of NLRP3 protein during follicular development in the mouse ovary. Positive staining in brown, counterstaining with heamatoxylin. (H, J) Negative control incubated with secondary antibody. Localisation of NLRP3 in (I) whole ovary of 16 weeks (wk) mice fed chow diet (CD), (K) primary follicles of 16 wk CD mice, (L) secondary follicles of 16 wk high fat diet (HFD) mice, and (M) preantral follicles of 16 wk CD mice. Staining was detected in granulosa and theca cells. Faint staining was observed in the oocytes of all stages of folliculogenesis. Arrows denote granulosa cells, arrow-heads denote theca cells, asterisks denote oocytes. The immunohistochemistry staining was confirmed by immunofluorescent localisation of NLRP3. Positive staining in orange, nuclear counterstaining with DAPI in blue. (N) Negative control 16 wk CD stained with polyclonal rabbit IgG. NLRP3 localised in (O) whole ovary, (P-Q) secondary follicles of 16 wk CD mice, (R-S) preantral follicles of 16 wk HFD mice, (T-U) antral follicles of 16 wk HFD mice. Inserts in top left corners represent magnifications of granulosa cells. Scale bars represent 20 or 100 μm.

### 4.2 Time-course activation of NLRP3-induced inflammasome in the ovary of DIO mice

In the following experiment, we tested the effect of short (4 wk) versus long term (16 wk) HFD treatment on NLRP3-induced inflammasome activation in the ovary of mice (**Figure 2A**). We used the DIO protocol previously validated, in which female mice were fed chow diet (CD) or HFD for 4 or 16 wk (n=8/group) (Wołodko et al. 2020). After the DIO protocol, we recorded the average body weight (BW) of 19.7 gram (g) in 4 wk CD group and 24.8 g in 4 wk HFD group, whereas the 16 wk CD presented on average BW of 22.6 g and 16 wk HFD 37.6 g BW (**Table 3**). After collection, ovaries were processed for mRNA and protein expression analysis. Real-time PCR analysis (n=6-8) revealed increased mRNA of *Nlrp3* after 4 wk HFD (**Figure 2B**, p=0.06), whereas *Il-1*β levels were increased after both 4 wk and 16 wk of HFD (**Figure 2B**, p<0.05). Regarding Western blotting analysis (n=8), we found the expression of NLRP3, CASP1 p45 and pro-IL-18 p24 were increased in 4 wk HFD group, compared to control group (**Figure 2C, D, and F**, p<0.05, respectively). However, opposite pattern was observed in 16 wk HFD, with downregulation of NLRP3 expression, the mature form of CASP1 p20 and both forms of IL-18 (p24 and p18) (**Figure 2 C, E, F and G**, p<0.05). Finally, we also confirmed IL-1β protein level was upregulated in 16 wk HFD after measurement by enzyme linked immunosorbent assay (ELISA) (n=6) (**Figure 2H**, p=0.082). Therefore, we presently confirmed that NLRP3, the pro-proteins IL-18 (p18, p24) and CASP1 (p45) despite being upregulated after 4 wk HFD treatment, after 16 wk HFD treatment a consistent downregulation of NLRP3 inflammasome, particularly NLRP3, CASP1 (p20) and both forms of IL-18, was seen. As a result, increased IL-1β protein after 16 wk HFD should be promoted independently from the NLRP3 inflammasome pathway (Lukens et al. 2014; Ranson et al. 2018).

**Figure 2.**
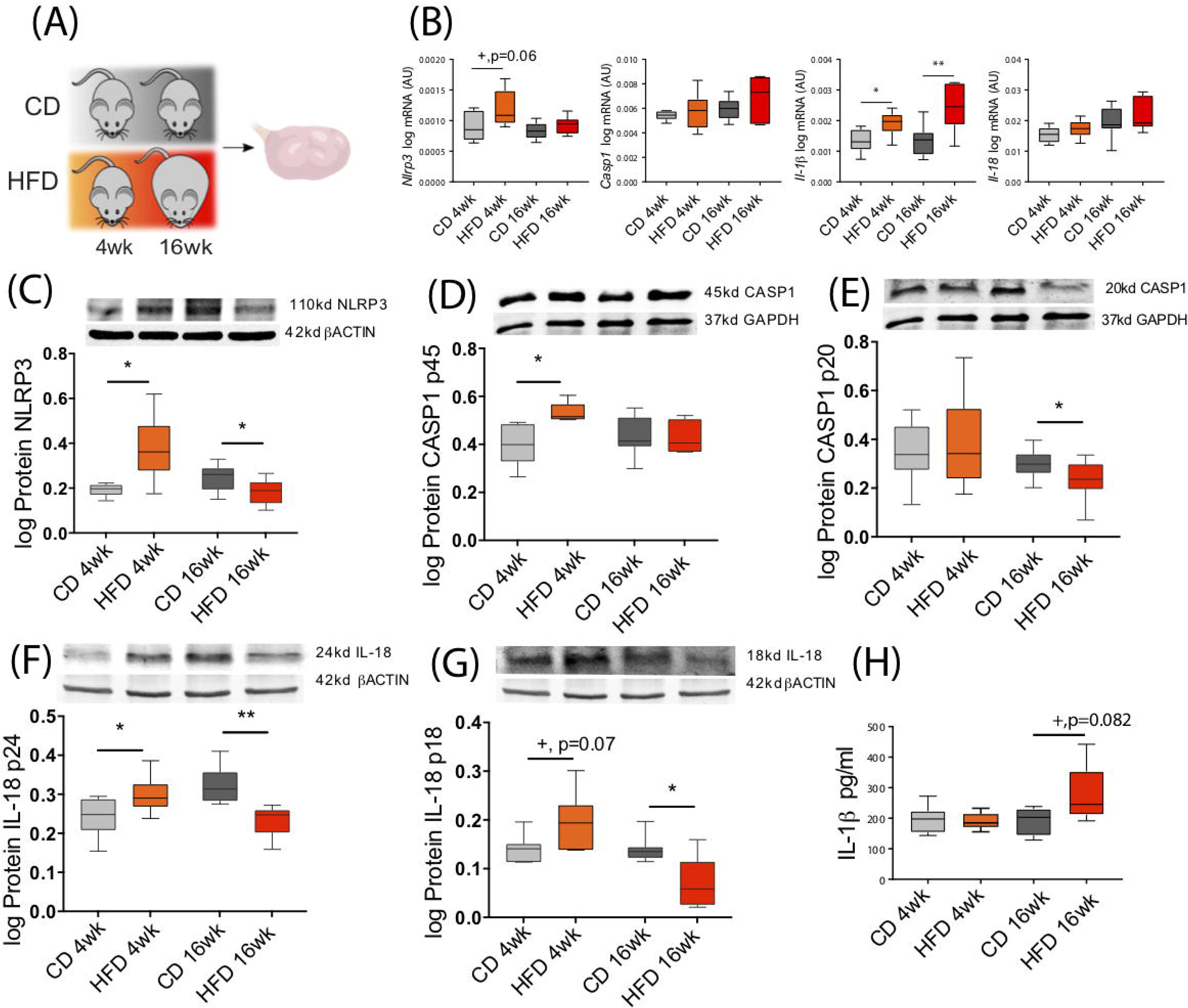
Diet induced-obesity changes NLRP3 expression in the ovary. (A) Experimental design: mice were fed either chow diet (CD) or high fat diet (HFD) for 4 weeks (wk) or 16 wk and ovaries were collected during dioestrus stage. Quantification of (B) Nod-Like Receptor Protein 3 (*Nlrp3*), caspase-1 (*Casp1*), interleukin-1β (*Il-1*β), and interleukin-18 (*Il-18*) mRNA by real-time PCR. Abundance of (C) NLRP3, (D) pro CASP1 p45, (E) CASP1 p20, (F) pro IL-18 p24, (G) IL-18 p18 protein measured by western blotting and (H) IL-1β protein measured by ELISA in ovarian extracts collected from DIO mice. mRNA level was normalized with ribosomal protein L37 (*Rpl37*) value and protein expression with β actin or glyceraldehyde 3-phosphate dehydrogenase (GAPDH) level. Bars represent mean ± SEM. Differences between control and treatment groups analysed with Mann–Whitney; n=6-9 for real-time PCR analysis, n=8 for immunoblots and n=6 for ELISA. Asterisks indicate significant differences (*p< 0.05; **p<0.01; +p=0.06; +p=0.07 or +p=0.082 – all values indicated).

**Table 3.**
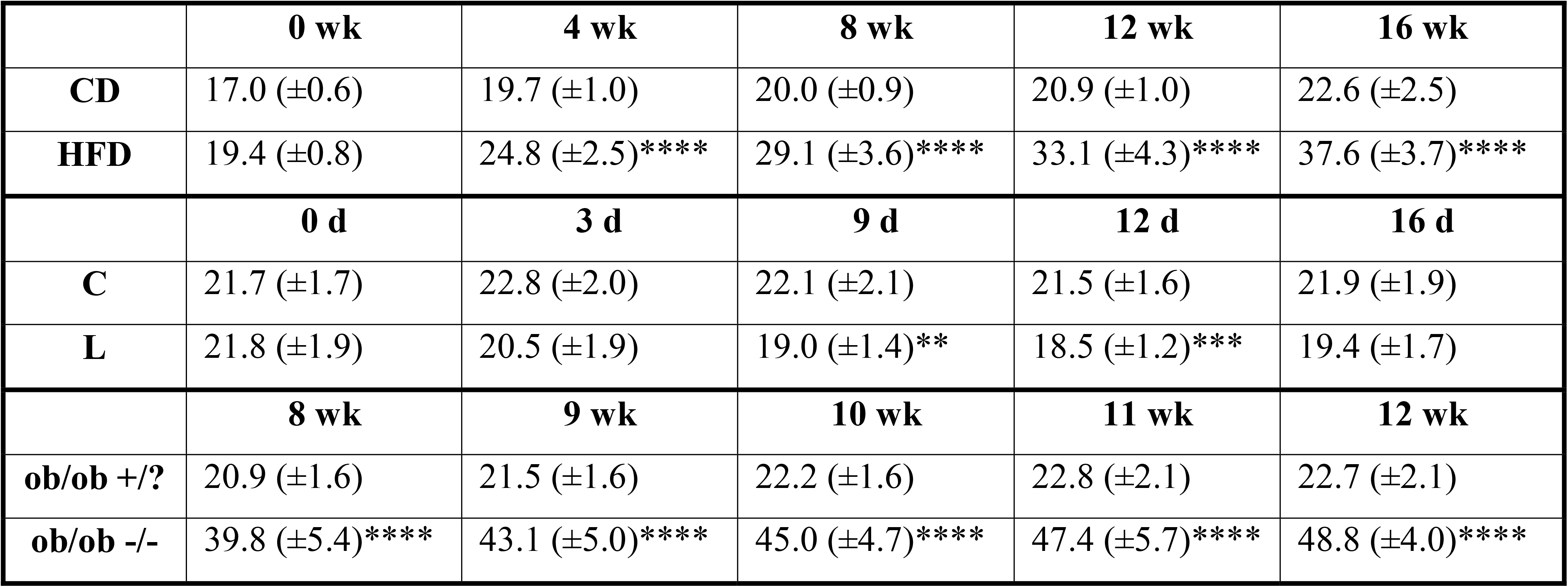
Body weight measurement of 3 mouse models. Diet induced obese (DIO) mice were fed chow diet (CD) or high fat diet (HFD); ii) pharmacologically hyperleptinemic mice were treated with saline (C) or leptin (L) for 16 days; iii) genetically obese mice lacking leptin (*ob/ob* -/-) and control group (*ob/ob* +/?). Statistical analysis between groups was carried out using T-test. Asterisks indicate significant differences (**p< 0.01; ***p< 0.001; ****p<0.0001).

### 4.3 Leptin signalling in the ovary drives activation of NLRP3 inflammasome during obesity progression

After temporally characterising the expression profile of NLRP3-induced inflammasome components in the ovary of DIO mice, we further interrogated whether activation of the NLRP3 inflammasome was regulated by leptin signalling. Indeed, leptin was previously shown to modulate NLRP3 expression *in vitro* (Fu et al. 2017). Furthermore, the expression signature of NLRP3 inflammasome components in the ovaries of DIO mice overlapped both tyrosine 985 of leptin receptor (Tyr985ObRb) and Janus kinase 2 (JAK2) phosphorylation profile, with the increase at 4 wk HFD treatment being followed by inhibition at 16 wk HFD and concomitant establishment of leptin resistance (Wołodko et al. 2020). Therefore, we analysed the levels of NLRP3 inflammasome components in the ovaries of a previously validated mouse model of pharmacological hyperleptinemia, which presented increased systemic levels of leptin and increased leptin signalling in the ovary without obesity (Wołodko et al. 2020), and a genetically obese mouse B6.Cg-Lepob/J (*ob/ob*), characterised by extreme obesity without leptin. In the pharmacological hyperleptinemic model, ten female B6 8 wk old mice, were treated with leptin intraperitoneally, twice a day for 16 days (16 L), whereas controls were administered saline (16 C) (Wołodko et al. 2020). Moreover, ten female *ob/ob* control (+/?) and ten females homozygous mutant (-/-), 8 wk old were kept on CD for 4 wk (**Figure 3A**). Ovaries from all groups were collected and processed for mRNA and protein expression analysis. Real-time PCR analysis (n=6-8/group) revealed an increase in *Nlrp3* and *Casp1* in 16 L, but decrease in *ob/ob* -/- mice (**Figure 3B**, p<0.05). Furthermore, mRNA of *Il-1*β was upregulated in 16 L (**Figure 3B**, p<0.05). Finally, the mRNA of *Il-18* was significantly downregulated in *ob/ob* -/- group (**Figure 3B**, p<0.05). With regard to protein expression, we found the 16 L group presented increased levels of NLRP3 (**Figure 3C**, p<0.05), whereas the opposite pattern was observed in *ob/ob* -/- mice, comparing to control groups (**Figure 3C**, p<0.05). Accordingly, both pro-peptides CASP1 (p45) and CASP1 (p20) showed increased levels in 16 L (**Figure 3D, E**; p=0.07 and p<0.05, respectively), nonetheless, no significant changes were found in the *ob/ob* model. Importantly, IL-1β protein measured by ELISA was increased in 16 L, but decreased in *ob/ob* -/- (**Figure 3F**, p<0.05). In this experiment we revealed the functional link between leptin signalling and NLRP3 inflammasome components regulation in the ovary, with leptin treatment inducing the activation of NLRP3 and CASP1 with subsequent secretion of IL-1β Furthermore, the absence of NLRP3 inflammasome activation in the ovary of *ob/ob* -/- confirms the preponderant role active leptin signalling exerts on NLRP3 inflammasome activation in the ovary.

**Figure 3.**
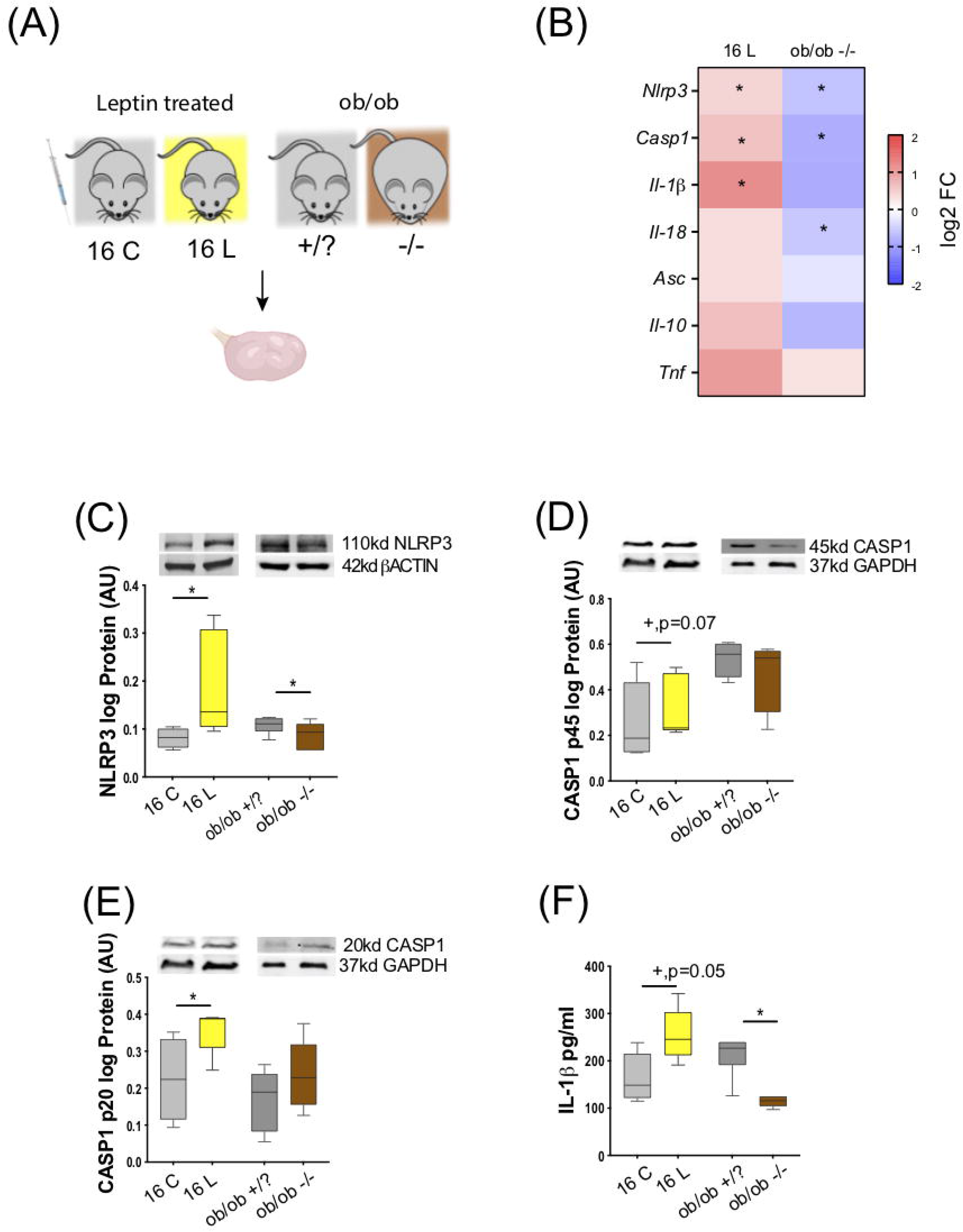
Leptin signalling in the ovary drive changes of NLRP3 during obesity. A) Experimental design: i) pharmacological hyperleptinemia model, mice were either injected with saline or 100µg of leptin (L) for 16 days; and ii) genetic obesity model, mice lacking leptin in circulation (*ob/ob* -/-) or control group (*ob/ob* +/?). B) Heatmap illustrating fold of change in expression of mRNA of genes Nod-Like Receptor Protein 3 (*Nlrp3*), caspase-1 (*Casp1*), interleukin-1β (*Il-1*β, interleukin-18 (*Il-18*), apoptosis-associated speck-like protein containing A CARD (*Asc*), interleukin-10 (*Il-10*) and tumour necrosis factor alpha (*Tnf*) in hyperleptinemia and *ob/ob* models determined by real-time PCR. The scale matches colours to log 2 fold change (log2_FC) values. Abundance of (C) NLRP3, (D) pro CASP1 p45, (E) CASP1 p20 measured by wersten blotting and (F) Il-1β quantified by EIA, in the mouse ovarian extracts. Level mRNA normalized with ribosomal protein L37 (*Rpl37*) value and protein expression with β actin or glyceraldehyde 3-phosphate dehydrogenase (GAPDH) . Bars represent mean ± SEM. Statistical analysis between groups was carried out using Mann–Whitney test. n=6-9 for real-time PCR analysis and n=4-8 for immunoblots. Asterisks indicate significant differences (*p< 0.05; + p=0.07).

### 4.4 Leptin promotes changes of NLRP3 inflammasome components gene expression in cumulus cells during early onset of obesity

In this experiment we examined whether the association previously observed between leptin signalling and NLRP3 inflammasome activation found in whole ovaries holds true at the cellular level, particularly for the somatic companions of the female gamete, the CCs. Indeed, the ovary is a very heterogeneous organ, with follicles in different developmental stages, and different somatic cells supporting oocyte development (Chang, Qiao, and Leung 2017). Therefore, we reanalysed our RNA sequencing (RNA-seq) datasets from CCs from the 4 wk HFD, 16 wk HFD and 16 L groups (Wołodko et al. 2020). Briefly, we collected approximately 50 CCs per animal, after superovulation, and RNA-seq libraries were generated using a Smart-seq2 oligo-dT method (**Figure 4A**; Wołodko et al. 2020). We started confirming the expression level of leptin and NLRP3 pathway components for 16 L and 4 wk HFD. Despite no changes in *Nlrp3* in CCs after 4 wk HFD, the gene was upregulated in 16 L (**Figure 4B**). Certainly, the low coverage of the samples (an average of 5.5 million reads) and the weak expression level of *Nlrp3* in CCs may account for the lack of changes in 4 wk HFD. Nonetheless, the consistent upregulation of various components of the NLRP3 inflammasome, like *Nlrp3* itself, or *Il-18, Casp1, Il-1*β and *Asc* in 16 L, is suggestive of the stimulatory effect of leptin on the expression of NLRP3 inflammasome genes also in CCs (**Figure 4B**). As previously shown, DESeq analysis revealed 997 differentially expressed genes (DEGs) in 4 wk HFD and 2026 DEGs in 16 L (Wołodko et al. 2020), in comparison to their control groups (p<0.05; Wołodko et al. 2020). In the present analysis, we overlapped the DEGs from 4 wk HFD and 16 L and identified seven genes either up- or downregulated in both conditions (**Figure 4C**). Subsequently, we integrated these 14 DEGs with the main components of NLRP3 and leptin signalling pathways (Wołodko et al. 2020) based on the correlation between their expression levels (p>0.90), obtaining five clusters, with one of them underscoring the gene interaction between *Casp1*, *phosphatase and tensin homolog (Pten)* and *signal transducer and activator of transcription 5a (Stat5a)*, as well as the link between *Socs3* and *Il-1β,* known as an important axis involved in the mediation of immune response (Chaves de Souza et al. 2013). Importantly, other genes were highlighted in the network, as *solute carrier family 22 member 15 (Slc22a15)*, a cell membrane transporter and metabolic gene (Nigam 2018), or *stress associated endoplasmic reticulum protein 1 (Serp1)* involved in protein unfolding and stress response (Yamaguchi et al. 1999). Indeed, metabolic performance in the preovulatory follicle is tightly regulated and involves the crosstalk between GC and oocyte (Wołodko et al. 2021). Also, the ER stress is a common feature observed in the ovaries of obese mothers (Robker, Wu, and Yang 2011) (**Figure 4D**). Finally, gene ontology analysis for the presented network revealed three main events, as negative regulation of glucose transport, positive regulation of cytokine biosynthesis and response to ATP (**Figure 4E**, p<0.05). Those certainly are key processes for oocyte maturation, as glucose metabolism in GC provides energy supplies for oocyte maturation (Wołodko et al. 2021). Furthermore, energy production through lipid oxidation and ATP production, is also fundamental for oocyte maturation (Wołodko et al. 2021). Finally, we plotted a subset of genes known to directly activate the NLRP3 inflammasome signalling pathway (Weber et al. 2020), particularly regarding the regulation of glutathione, major mediator of NLRP3 signalling (Hughes et al. 2019), as well as other genes involved in the pathway regulation (Barlan et al. 2011; Billon et al. 2019; Guglielmo et al. 2017; Y. He et al. 2016; Hughes et al. 2019; Hughes and O’Neill 2018; Iyer et al. 2013; Jo et al. 2016; Kim et al. 2015; Li et al. 2016; Martine et al. 2019; Mitoma et al. 2013; Palazón-Riquelme et al. 2018; Shuvarikov et al. 2018; X. Wang et al. 2014; Wolf et al. 2016; T. Zhang et al. 2021; Zhou et al. 2010), and confirmed the similarities between 16 L and 4 wk HFD for those gene lists, in opposition to 16 wk HFD (**Figure 4F**). Thus, as presently shown, systemic administration of leptin activated genes from the NLRP3 inflammasome pathway in CCs, corroborating once more the functional link between leptin signalling and NLRP3 inflammasome activation in CCs of DIO mice.

**Figure 4.**
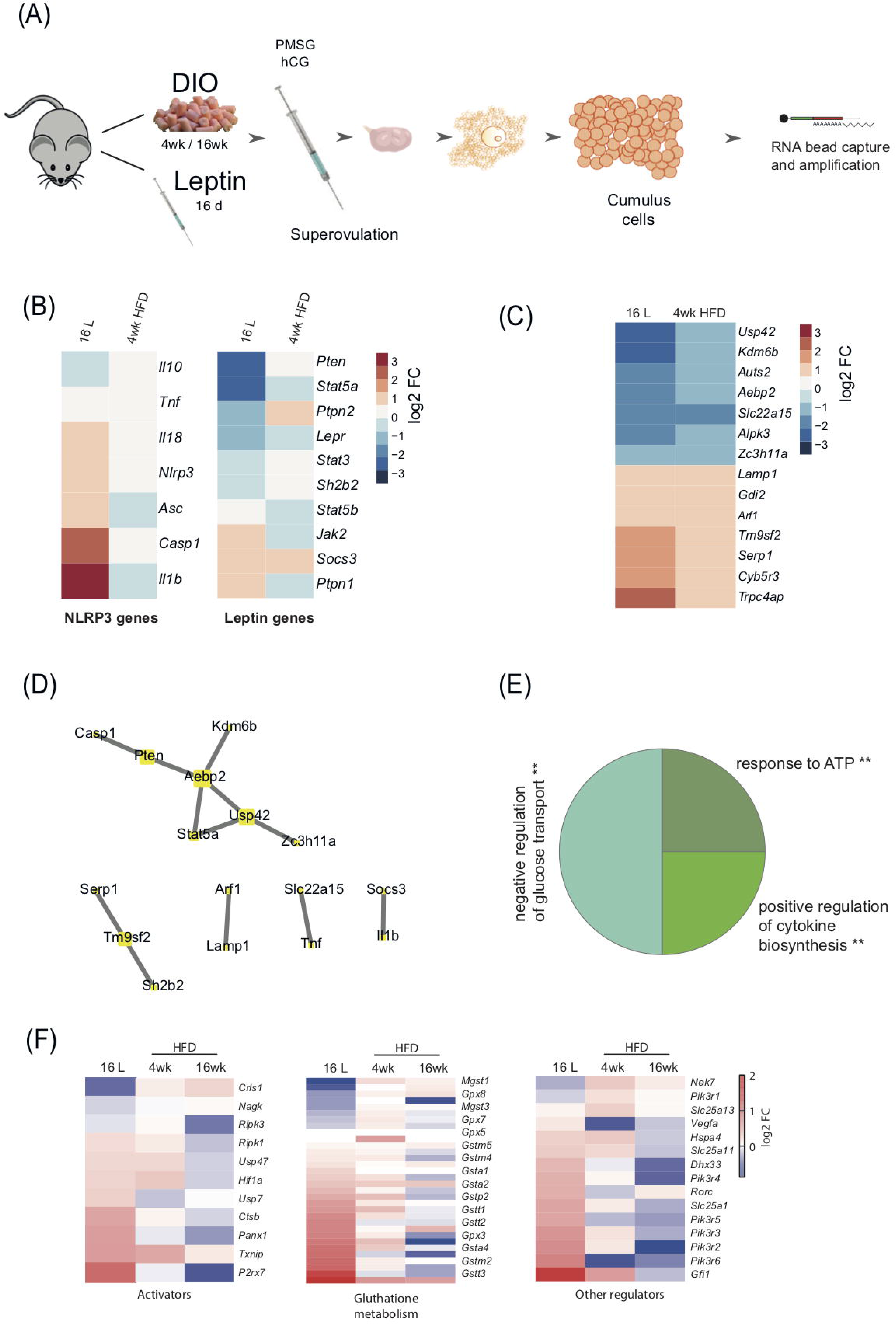
Cumulus cells transcriptome analysis from diet-induced obese and pharmacologically hyperleptinemic mice reveals leptin-mediated regulation of NLRP3 inflammasome genes. A) Experimental design: RNA-Seq analysis of differentially expressed genes in cumulus cells collected from mice: i) with diet-induced obesity (DIO) fed chow diet (CD) or high fat diet (HFD) for 4 weeks (wk) or 16 wk ii) with pharmacologically hyperleptinemic (LEPT) injected with saline (C) or 100µg of leptin (L) for 16 days. B) Heatmap illustrating expression of genes from leptin signalling pathway and NLRP3 inflammasome in LEPT and 4 wk HFD normalized by control group fed CD or injected with saline, respectively. Downregulated genes in blue, upregulated genes in orange. The scale on the right matches colours to log 2 fold change (log2 FC) values. (C) Heatmap showing significant changes in major constitutive genes in both conditions leptin and 4 wk HFD. (D) Five main clusters in the network representing strong interaction between selected genes described in B and genes presented in C. (E) Pie chart that displays the main three gene ontology terms that were significantly enriched in cumulus cells in both conditions LEPT and 4 wk HFD. Gene ontology analysis performed with Gene Ontology Enrichment Analysis and Visualisation Tool. (F) Heatmaps showing conserved genes involved in NLRP3 inflammasome activation, glutathione metabolism and other regulations. log2 FC of reads per million (RPM).

### 4.5 Time-course activation of NLRP3-induced inflammasome in liver of DIO mice

Different organs can uniquely adapt to systemic insults like obesity (Smith et al. 2018). Therefore, in the last experiment was asked to what extent mounting inflammatory response through NLRP3 inflammasome activation in the ovaries of DIO mice precede other metabolic organs like the liver. We analysed the expression profile of NLRP3 inflammasome genes in the liver of DIO mice, besides testing once more the functional link between leptin signalling and activation of NLRP3 inflammasome at hepatic level, using pharmacological hyperleptinemic and *ob/ob* mouse models. Liver samples were collected from DIO, leptin treated and *ob/ob* female mice, for mRNA transcription and protein expression analysis (**Figure 5A**). Real-time PCR analysis (n=5-7/group) showed no significant changes in expression of all inflammasome components, except for the increase of *Nlrp3* and *Il-1*β mRNA in 16 wk HFD, but downregulation of *Nlrp3* in *ob/ob*-/- (**Figure 5B**, p<0.05). Furthermore, protein analysis determined by Western blotting (n=4-8/group) showed an increase in protein levels of NLRP3, CASP1 (p20), and IL-18 (p18) after 16 wk HFD treatment, comparing to control (**Figure 5C, E and G**, p=0.07, p<0.05). Regarding the pro-peptide of CASP1 (p45), its protein was upregulated in 16 L, but downregulated in *ob/ob* -/-, comparing to controls (**Figure 5D**, p<0.05 and p<0.01, respectively). Finally, CASP1 (p20) and IL-18 (p18) proteins were decreased in *ob/ob*-/- (**Figure 5E, G**, p<0.05 and p=0.08, respectively). No significant changes were observed for pro-IL-18 (p24). The present results on our analysis in the liver indicate a site dependent NLRP3 inflammasome regulation throughout obesity, since overexpression of NLRP3, CASP1 (p20) and IL-18 (p18) took place only at 16 wk of DIO. Differences in NLRP3 inflammasome profile between liver and ovary certainly relay on the intrinsic immunological complexity the liver presents. The liver, in opposition to the ovary, is constantly exposed to proinflammatory mediators, having developed the ability to tightly control inflammation (Robinson, Harmon, and O’Farrelly 2016). Another important observation was the downregulation of NLRP3 and CASP1 (p20) in livers from *ob/ob* -/- mice. Other studies corroborated these observations (Negrin et al. 2014), and despite all intricacies of NLRP3 inflammasome regulation, leptin seems to directly modulate NLRP3 inflammasome activation at hepatic level. Hence, we confirmed the latency of NLRP3 inflammasome activation in the liver of DIO female mice, which showed signs of upregulation only after 16 wk HFD treatment. Furthermore, we have confirmed the functional link between leptin and NLRP3 inflammasome activation in the liver.

**Figure 5.**
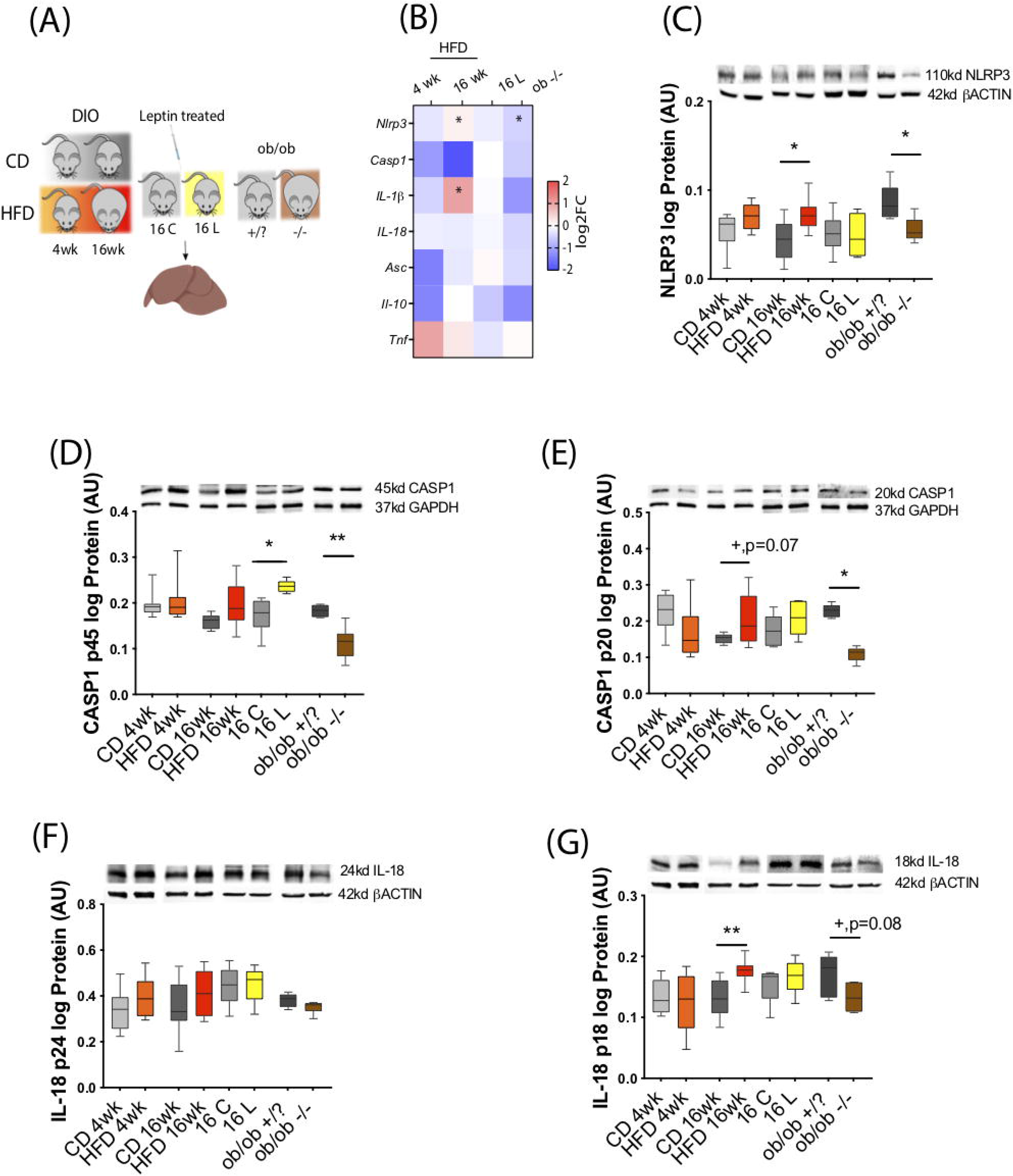
NLRP3 activity in the liver of diet-induced obese, hyperleptinemic and genetically obese mice. (A)Experimental design: i) diet induced obese (DIO) mice were fed chow diet (CD) or high fat diet (HFD) for 4 weeks (wk) or 16 wk; ii) pharmacologically hyperleptinemic mice were treated with saline (C) or 100µg of leptin (L) for 16 days; iii) genetically obese mice lacking leptin (*ob/ob* -/-) and control group (*ob/ob* +/?). (B) Heatmap illustrating fold change in expression of mRNA of genes Nod-Like Receptor Protein 3 (*Nlrp3*), caspase-1 (*Casp1*), interleukin-1β (*Il-1*β, and interleukin-18 (*Il-18*), apoptosis-associated speck-like protein containing a CARD (*Asc*), interleukin-10 (*Il-10*) and tumour necrosis factor alpha (*Tnf*) in DIO, hyperleptinemia and *ob/ob* models in comparison to respective controls, determined by real-time PCR. The scale on the right matches colours to log2 fold change (log2 FC) values. Data normalised to mRNA expression of ribosomal protein L37 (*Rpl37*). Abundance of (C) NLRP3, (D) pro CASP1 p45, (E) CASP1 p20, (F) pro IL-18 p24 in mouse liver of DIO, hyperleptinemic and *ob/ob* mice measured in western blotting analysis. Protein normalised with β-actin or glyceraldehyde 3-phosphate dehydrogenase (GAPDH) level. Bars represent mean ± SEM. Statistical analysis between groups was carried out using Mann–Whitney test; n=6-9 for real-time PCR analysis and n=5-8 for immunoblots. Asterisks indicate significant differences (*p< 0.05; **p<0.01).

## 5 Discussion

The present study gives the first characterisation of NLRP3 induced inflammasome activation in the ovaries of DIO mice. Maternal obesity has been largely associated with increased ovarian inflammation (J. Nteeba et al. 2013; Jackson Nteeba, Ganesan, and Keating 2014; Robker, Wu, and Yang 2011; Ruebel et al. 2017; Snider and Wood 2019), being a better knowledge of its pathogenesis of undeniable value for our understanding of ovarian failure and infertility during obesity. We firstly confirmed the effects of cyclicity on NLRP3 inflammasome activation in the ovaries of lean mice, observing the upregulation of NLRP3 inflammasome components in D. Subsequently, we temporally characterised the expression profile of NLRP3 inflammasome components in the ovary, throughout obesity progression. Indeed, the rapid upregulation of NLRP3 protein in early obesity (after 4 wk HFD treatment), was followed by a consistent downregulation of NLRP3 inflammasome components, as NLRP3 and CASP1, in late obesity (after 16 wk HFD). Importantly, using either a pharmacological hyperleptinemic and a genetic obese *ob/ob* mouse, we not only evidenced the functional link between levels of leptin signalling and NLRP3 activation in whole ovaries, but also the role of leptin on *Nlrp3*, *Il-18 and Il-1*β gene expression upregulation in CCs from ovulated follicles. Finally, after analysing the NLRP3 inflammasome expression pattern in the liver, we confirmed NLRP3 and CASP1 overexpression took place exclusively after 16 wk HFD treatment, suggesting a delayed activation of NLRP3 inflammasome activation in comparison with the ovary. Hence, these results suggest a greater vulnerability of the ovaries in general, and the gamete in particular, to the energetic surplus females face under obesogenic conditions.

A recent study by Zhang and colleagues evidenced for the first time NLRP3 expression in various cellular components like GC, TC and oocytes of mouse ovaries (Z. Zhang, Wang, and Zhang 2019). We presently confirmed not only the similar pattern of cellular expression for NLRP3, but also the upregulation of NLRP3 inflammasome components during D. These findings corroborate previous results suggesting the involvement of NLRP3 in inflammation during ovulation in mice (Z. Zhang, Wang, and Zhang 2019). Furthermore, NLRP3 proinflammatory role in ovarian function starts getting noticed not only under physiological context (Z. Zhang, Wang, and Zhang 2019), but also as an important mediator of ovarian pathology during ageing (Navarro-Pando et al. 2021). Certainly, our hypothesis of NLRP3 inflammasome involvement in inflammatory response in the ovary of obese mothers seems to be supported also by earlier reports showing NLRP3 inflammasome activation during development and treatment of PCOS (Guo et al. 2020; F. Wang et al. 2017).

The NLRP3 inflammasome is a critical component of innate immunity, frequently associated with human disease (Y. He, Hara, and Núñez 2016). Our results evidencing NLRP3 inflammasome activation in the ovaries of 4 wk HFD treatment are in line with previous reports showing the accumulation of proinflammatory mediators in the ovary after short term (6 wk HFD) dietary protocols (Shen, Xu, and Li 2021). Indeed, we presently observed the upregulation of IL-18 protein, as well as increased mRNA of *Tnf* (data not shown), after 4 wk HFD underscoring the mounting inflammatory response. Nonetheless, maintenance of inflammation in the ovaries in long term DIO mice (after 16 wk HFD) seems to be mediated independently from NLRP3 inflammasome pathway, as IL-18 and CASP1 mature proteins were significantly downregulated at this time point. Undeniably, increased levels of IL-1β in the ovaries of 16 wk HFD mice confirm the proinflammatory state in the ovaries of DIO mice after 16 wk HFD, supported by several studies in mice showing the abundancy of inflammatory markers like Il-1β, *Il-6* and *Tnf*α and in the ovary of long term DIO mice fed for 24 wk HFD (J. Nteeba et al. 2013). As a result, the increased levels of IL-1β at 16 wk HFD confirm the existence of alternative pathways to NLRP3 inflammasome, mediating IL-1β upregulation (Donado et al. 2020; Jain et al. 2020; Pyrillou, Burzynski, and Clarke 2020; Schmidt and Lenz 2012; Zhu and Kanneganti 2017). To this extent, we have recently shown the temporal pattern of expression of proinflammatory genes in CCs from DIO mice (Wołodko et al. 2020). In early obesity, inflammatory cues in CCs were mediated by cellular response to stress through upregulation of genes like *DEAD-box helicase 5 (Ddx5), hypoxia inducible factor 1 subunit alpha (Hif1a), ADAM metallopeptidase domain 9 (Adam9*). Indeed, mediators of stress response as ROS are known to prime the NLRP3 inflammasome (Gurung et al. 2014). Subsequently, in late obesity, we saw the overexpression of genes involved in anatomical structural morphogenesis, as *C-C motif chemokine ligand 7* (*Ccl7*), an important chemoattractant of leukocytes (Menten et al. 1999), and also known to interact with matrix metalloproteinases (MMPs) (Liu et al. 2018), or complement C3a receptor 1 (*C3ar1*), a complement component known to mediate neutrophil mobilisation (Brennan et al. 2019) and lately described as a marker of PCOS progression (D. He et al. 2020). Therefore, these data suggest important temporal dynamics on the regulation of the inflammatory response in the ovary throughout obesity progression, with NLRP3 inflammasome playing a critical role mostly in the initiation of inflammation in the ovaries in early obesity. Conversely, in late obesity, immune mediated response in the ovary progresses to infiltration of immune cells and structural reorganisation, independently from the activation of NLRP3 inflammasome.

Our study also sheds light on the important crosstalk between leptin signalling and inflammasome NLRP3 activation in the ovary of DIO mice. As reviewed by Wani and co-workers, numerous factors were shown to activate NLRP3 inflammasome during obesity, such as cellular metabolites, carbohydrates or lipids (Wani et al. 2021). Nonetheless, leptin, a conserved proinflammatory cytokine (Iikuni et al. 2008; La Cava 2017), was recently shown to upregulated NLRP3 components *in vitro* (Fu et al. 2017). Thus, in order to test the hypothesis whether repression of NLRP3 inflammasome activation in ovaries of 16 wk HFD mice was due to the establishment of leptin resistance (Wołodko et al. 2020), we studied NLRP3 inflammasome activation in the ovaries of pharmacological hyperleptinemic and *ob/ob* mice. Strikingly, we observed a consistent upregulation of NLRP3 inflammasome genes and accumulation of IL-1β protein in ovaries of leptin treated mice, in opposition to *ob/ob* -/- mice which evidenced consistent downregulation of NLRP3 and IL-1β proteins. Furthermore, reduced levels of NLRP3 were also observed in *ob/ob* -/- mouse peritoneal macrophages treated with LPS and nigericin in comparison to wild type mice (Yang et al. 2021), what certainly underlines the preponderant role of leptin on NLRP3 inflammasome regulation. Therefore, our results invite us to suggest the activation of NLRP3 inflammasome in the ovary of DIO mice is mediated by leptin signalling. In early obesity (4 wk HFD treatment) leptin actively signals through receptor b (ObRb) in the ovary (Wołodko et al. 2020), with the overexpression of NLRP3 inflammasome components; nonetheless, in late obesity (16 wk HFD treatment) after the establishment of leptin resistance in the organ (Wołodko et al. 2020), expression of NLRP3 inflammasome is drastically repressed.

Next, we reanalysed our datasets on global gene expression in CCs collected from pharmacological hyperleptinemic and DIO mice, in order to test the association between leptin signalling and activation of NLRP3 inflammasome in the somatic companions of the oocyte. Importantly, CCs are known as faithful indicators of intrafollicular environment (Wołodko et al. 2021) and their transcriptome has been used to predict oocyte and embryo quality (Uyar, Torrealday, and Seli 2013). Despite no changes in 4 wk HFD, we confirmed the overexpression of NLRP3 inflammasome genes in CCs from 16 L. Consequently, we interrogated whether DEGs overlapping both 16 L and 4 wk HFD treatment could interacted with NLRP3 inflammasome genes. Indeed gene ontology for associated genes revealed key terms for oocyte maturation, as regulation of glucose transport, response to ATP and regulation of cytokine biosynthesis. Metabolic regulation in preovulatory follicles appears to control major steps for maturation of female gamete, as meiosis resumption, chromatin condensation and cytoplasm maturation (Wołodko et al. 2021). For instance, glucose, which is mostly metabolised in CCs (Sanfins, Rodrigues, and Albertini 2018) was shown to be key for oocyte competence (Wilding et al. 2009), as well as reduced ATP content in oocytes was linked to failure in fertilisation, arrested division and abnormal embryonic development (J. Zhao and Li 2012). Furthermore, the aforementioned involvement of NLRP3 inflammasome in ovulation (Z. Zhang, Wang, and Zhang 2019) can be considered amongst the regulation of the cytokine milieu locally produced in CCs. Given leptin direct and indirect role in ovulation (Wołodko et al. 2021), failure in leptin signalling and NLRP3 inflammasome activation in late obesity can account for increased anovulatory rates in obese mothers (Hou et al. 2016; Wu et al. 2010). Finally, absence of changes in 4 wk HFD in NLRP3 inflammasome genes can be ascribed to low coverage of our reduced-cell libraries and also low levels of gene expression. Indeed, the present RNA-seq protocol used as little as 50 cells per mouse, which has limitations while analysing weakly expressed genes. Collectively, our results indicate leptin and NLRP3 inflammasome crosstalk in CCs can interfere with major steps regulating oocyte maturation and early embryo development.

In the last experiment we confirmed the liver, in sharp contrast to the ovary, activated NLRP3 inflammasome later in time during DIO protocol (after 16 wk HFD treatment) in mice. Temporal differences in inflammatory response regulation between both organs certainly rely on contrasting exposition to exogenous pathogens. The liver is an organ constantly exposed to proinflammatory mediators from dietary and commensal bacterial products (Robinson, Harmon, and O’Farrelly 2016). Thus, the hepatic immune system is constantly in contact with altered metabolic activity and regular exposition to microbial products, which results in persistent and tightly regulated inflammatory response (Robinson, Harmon, and O’Farrelly 2016). On the contrary, the ovary is not only a highly immunogenic organ constantly secreting large amounts of cytokines and immune mediators (Piccinni et al. 2021), but also more prone to rapidly mounting proinflammatory response during obesity. Indeed, the inability of the ovary to control inflammation and exacerbated cytokine production certainly ascribes for the great vulnerability the female gamete presents to maternal obesity even at earlier stages. Thus, our results expose the increased ovarian vulnerability to maternal obesity, with a rapid mounting inflammation which affects the gamete and impairs fertilisation.

In summary, our work evidences the major role leptin signalling exerts on NLRP3 inflammasome activity in the ovary of obese mice. Noteworthy, failure in ovarian leptin signalling was associated with repression in NLRP3 activity, but not decreased inflammation and levels of IL-1β. Moreover, NLRP3 inflammasome activation in the ovary precedes liver response during obesity progression suggesting the greater vulnerability the ovary in general, and gamete in particular, to the energetic surplus during maternal obesity.

## Author Contributions

MA did data acquisition, analysis and interpretation of the data, writing the manuscript; KW did data acquisition and analysis and revised and edited the manuscript; JO did immunohistochemistry staining; JCF conducted data analysis and interpretation of data, revising the manuscript; DM did immunohistochemistry staining; GK revised and edited the manuscript; AG conceptualised and designed the study, acquired the funding, participated in data acquisition, analysis and interpretation, and wrote and edited the manuscript.

## Acknowledgments

We would like to thank Dr Leslie Paul Kozak and Dr Magdalena Jura for their support with the validation and characterisation of the mouse obese phenotype; and Dr Krzysztof Witek for his support with the imaging and confocal microscopy.

## Data Availability Statement

The raw data supporting the conclusions of this article were made available within the publication Wołodko et al 2020.

